# Mechanical Centrosome Fracturing during Cell Navigation

**DOI:** 10.1101/2024.10.01.615992

**Authors:** Madeleine T. Schmitt, Mauricio J.A. Ruiz-Fernandez, Kasia Stefanowski, Janina Kroll, Robert Hauschild, Jack Merrin, Thomas Penz, Eva Kiermaier, Christoph Bock, Tobias Straub, Jörg Renkawitz

**Affiliations:** Biomedical Center, Walter Brendel Center of Experimental Medicine, Institute of Cardiovascular Physiology and Pathophysiology, Klinikum der Universität, Ludwig Maximilians Universität München; Munich, Germany; Institute of Science and Technology Austria; Klosterneuburg, Austria; CeMM Research Center for Molecular Medicine of the Austrian Academy of Sciences; Vienna, Austria; Life and Medical Sciences (LIMES) Institute, Immune and Tumor Biology, University of Bonn; Bonn, Germany; Medical University of Vienna, Institute of Artificial Intelligence, Center for Medical Data Science, Vienna, Austria; Bioinformatics Unit, Biomedical Center, Faculty of Medicine, Ludwig Maximilians Universität München; Munich, Germany

## Abstract

The centrosome is the primary microtubule orchestrator in most eukaryotic cells, nucleating and anchoring microtubules that grow radially and exert forces on cargos. At the same time, mechanical stresses from the microenvironment and cellular shape changes compress and bend microtubules. Yet, centrosomes are membrane-less organelles, raising the question of how centrosomes withstand mechanical forces. Here we discover that centrosomes in non-dividing cells can mechanically fracture. We reveal that centrosomes experience mechanical deformations during microenvironmental confinement and navigational pathfinding by motile cells. Coherence of the centrosome is maintained by Dyrk3, preventing fracturing by mechanical forces. Centrosome fracturing impedes cellular function by generating coexisting microtubule organizing centers that compete during path navigation and thereby cause cellular entanglement in the microenvironment. Our findings show that non-dividing cells actively maintain the integrity of the centrosome to withstand mechanical forces. Given that almost all cells in multicellular organism experience forces, these results suggest that centrosome stability preservation is fundamental during development, tissue maintenance, immunology, and disease.

**One-Sentence Summary:** Pathfinding during cellular motility causes centrosome breakage counteracted by Dyrk3 activity.

## Main Text

Cells inside multicellular organisms experience mechanical forces that originate from intracellular cytoskeletal dynamics and coupling to the extracellular microenvironment, such as to neighboring cells and extracellular matrix (*1*). At the same time, cells have to endure these forces to prevent damage and to preserve functionality, including the maintenance of nuclear integrity when they squeeze through narrow gaps (*2, 3*). The microtubule cytoskeleton is one of the major sources of intracellular forces, moving cargos like vesicles during interphase and chromosomes during mitosis (*4*). Further, mechanical stresses from the microenvironment and from cellular shape changes exert forces on the microtubule cytoskeleton, which typically spans through the entire cell, resulting in microtubule compression or bending (*5, 6*). In many cells, the centrosome functions as the major nucleator and anchor of microtubules (*7*), which suggests that centrosomes evolved an architectural composition primed to withstand forces. Centrosomes are membrane-less organelles, composed of a pair of two centrioles that are connected by non-covalent linker proteins, and a surrounding proteinaceous matrix (*4*). While the membrane-less property of centrosomes is critical for their precisely timed separation during cell division, it raises the question how the mechanical integrity of the centrosome is maintained in non-dividing cells while experiencing forces.

As a model system for cells that experience forces (*8*), we live-imaged highly motile mouse dendritic cells (DCs) that are terminally differentiated (*9, 10*) and centrally nucleate microtubules only from the centrosome (*11*). To visualize the centrosome, we employed DCs that express centrin-2 (CETN2)-GFP (*10*) as a marker for the pair of two centrioles within one centrosome. The two centrioles remained at close proximity with minor distance fluctuations when cells migrated persistently along unidirectional paths along a chemotactic gradient (Fig. 1, A and B, and movie S1), in accordance with centrosome cohesion in interphase cells (*12*). Even during minor directional changes along wider straight paths, the centriole pair moved in a coordinated and synchronous manner (fig. S1, A and B). Yet, to our surprise, when the cells encountered path junctions, we observed that the distance between the two paired centrioles increased considerably, indicative for stretching deformations (Fig. 1, C and D, and movie S1). Centrosome stretching was transient, as the initial centriolar distance restored during cellular movement after productive path decision (Fig. 1E, and movie S1). These results suggested that centrosomes are subject to mechanical deformations, in line with historical observations of transient centriole separation during the adhesive spreading of neutrophils on surfaces (*13*).

**Fig 1.**
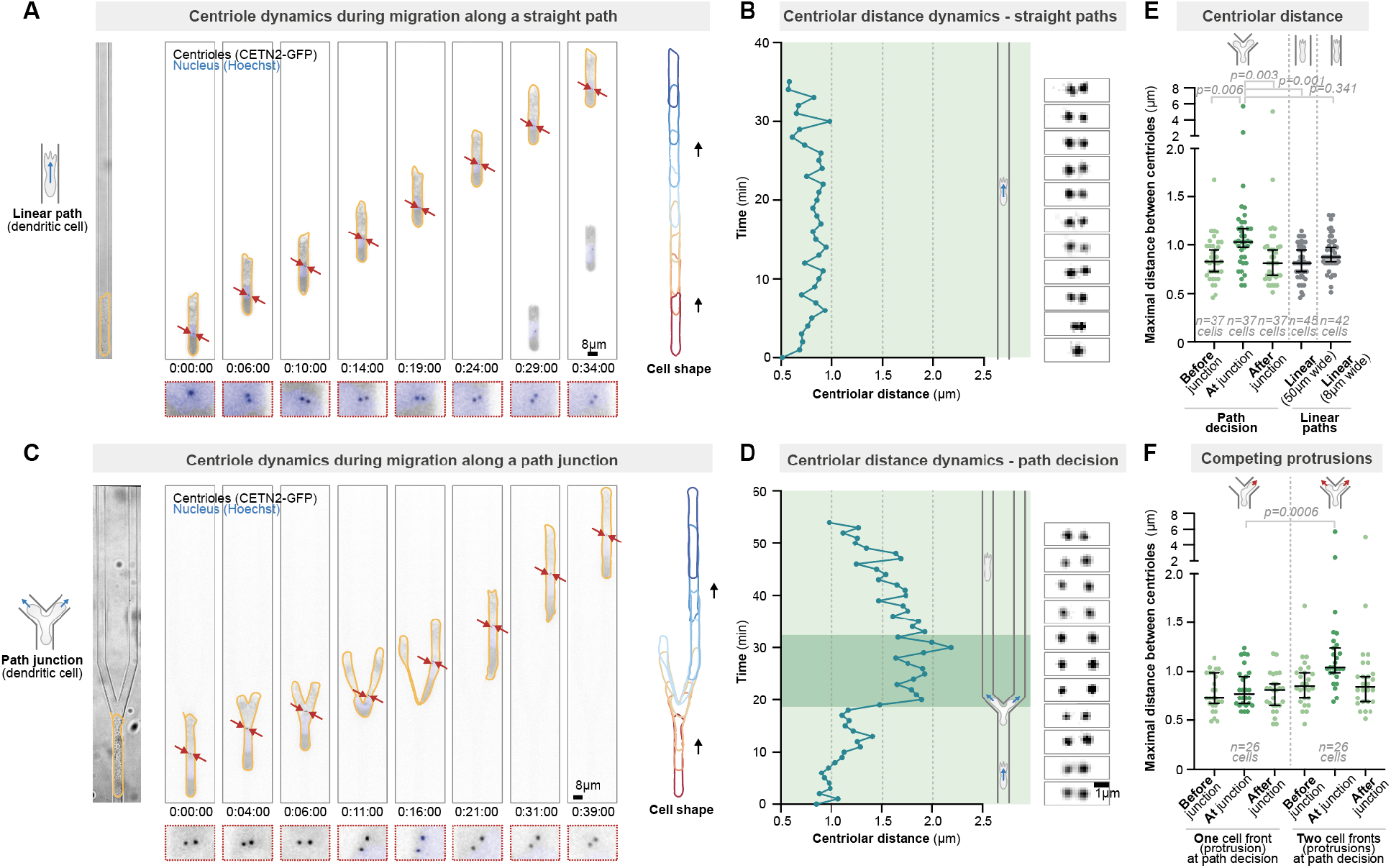
Mechanical centrosome deformations in motile cells. **>(A)** Representative CETN2-GFP (centriole pair: black; enlargement in red dashed boxes) expressing dendritic cell (DC) stained with Hoechst (nucleus; blue) migrating along a unidirectional straight path (linear microchannel). **(B)** Centriolar distance dynamics during migration along a unidirectional straight path; note the stable proximity of the individual centrioles. **(C)** As in (A), but migration through a path junction. Note the two cell fronts exploring the alternative paths. **(D)** Centriolar distance dynamics during a path decision; note the transiently increased distance between the individual centrioles. **(E)** Quantification of centriolar distances before, during, and after migration through path junctions, as well as during migration along narrow and wide linear paths. **(F)** Quantification of centriolar distances of cells that pass the junction either with two simultaneous explorative protrusions or DCs that immediately decide for one path alternative with one protrusion. Time is indicated as h:min:s.

Migrating DCs typically extend simultaneous cell fronts (protrusions) into alternative paths, a characteristic feature of motile cells to explore their microenvironment (*14, 15*). Thus, we compared centrosome deformations in DCs that either explored their path with two competing protrusions or immediately decided for one of the path alternatives with a single protrusion. DCs that migrated with two competing protrusions deformed their centrosome transiently as the two protrusions explored the alternative paths (Fig. 1, C, D, and F). Yet, DCs that migrated immediately with a singular protrusion into one path without exploring the alternative path, did not stretch the centrosome (Fig. 1F, and fig. S1, C and D). These findings show that forces from competing cellular protrusions are able to mechanically deform the centrosome. As centrosome stretching only rarely resulted in distant centriole separation (Fig. 1F), these data further suggested that centrosomes are equipped with mechanisms to actively withstand mechanical forces.

To screen for mechanisms that enable centrosomes to maintain their integrity during mechanical deformations, we established transcriptomics of migrating DCs in collagen networks of lower and higher complexity (fig. S2A), based on the rational that motile cells sample more complex environments with higher numbers of competing protrusions. Cluster analysis of differentially regulated genes revealed transcriptional adaptation to migration in more complex matrices (fig. S2B), including genes that are upregulated with increasing collagen complexity (fig. S2C). Given that the centrosome has features of membrane-less biocondensates (*16*–*18*), we were particularly interested in observing upregulation of Dyrk3 (fig. S2D), a protein kinase that has been shown to function as a regulator of membrane-less organelles (*19*), including its activity at the centrosome for cell cycle progression into mitosis (*20*). To test the role of Dyrk3 for locomotion of non-dividing interphase cells, we first investigated DC migration through collagen matrices in the presence of GSK-626616, a well-described small-compound inhibitor of Dyrk3 (*19, 20*). DCs showed reduced migration velocities during navigation through the collagen matrices (Fig. 2A, fig. S2E, and movie S2), which we also observed in the presence of harmine (Fig. 2B, and fig. S2F), an additional small-compound inhibitor of Dyrk-family proteins. Similarly, when we investigated human Jurkat T cells as another cellular model for fast cell migration, we observed reduced migration velocities and accumulated distances, but comparable directionality towards a chemokine source, in the presence of GSK-626616 (Fig. 2, C and E, fig. S2G, and movie S2) as well as upon expression of a dominant-negative kinase-dead point mutant of Dyrk3 (EGFP-DYRK3 K218M) (*20*) (Fig. 2, D and E, and movie S2). On the basis of these data, we conclude that functional Dyrk3 is required for efficient cell migration.

**Fig 2.**
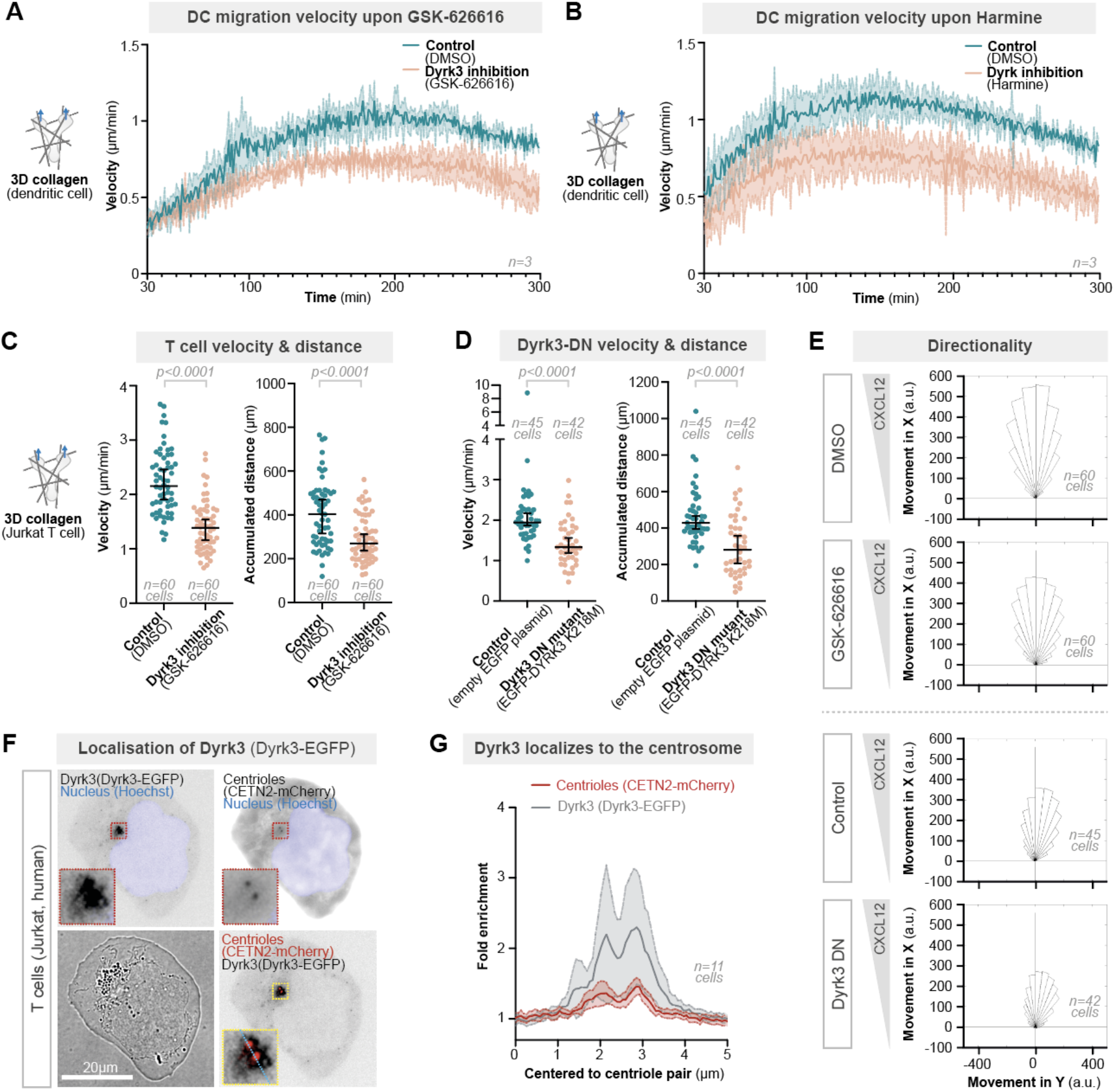
Cell motility requires functional Dyrk3. **(A)** Dendritic cell migration in three-dimensional (3D) collagen matrices (1.7 mg/ml) along a CCL19 chemokine gradient in the presence of 5μM GSK-626616 or DMSO (control). **(B)** As in A), but in the presence of 50μM Harmine. **(C)** Velocity and migrated distance of Jurkat T cells migrating in 3D collagen matrices (1.3 mg/ml) along a CXCL12 chemokine gradient in the presence of 5μM GSK-626616 or DMSO (control). **(D)** As in C), but comparing cells that express a dominant-negative (DN) EGFP-Dyrk3 K218 mutant or the corresponding empty EGFP plasmid. **(E)** Directionality along a chemotactic gradient (CXCL12) of Jurkat T cells upon rendering Dyrk3 non-functional. **(F)** Localisation of EGFP-tagged WT Dyrk3 and CETN2-mCherry in Jurkat T cells. **(G)** Quantification of (F) by measuring the fluorescent intensity along a 5μm line (blue dotted line in F) centered to the centriole pair.

As Dyrk3 (EGFP-DYRK3) localized primarily to the centrosome of motile cells (Fig. 2, F and G), we speculated that Dyrk3 might be critical for centrosome stability in the presence of motility forces. To test this hypothesis, we imaged CETN2-GFP expressing DCs during navigation through path junctions. Before the path junction, the centrioles typically located in close proximity and moved synchronously (Fig. 3, A and B). Yet, once the cells encountered multiple path options, where they typically explored most of the available paths at the same time, the pair of centrioles frequently broke into far-distantly located and individually moving centrioles when Dyrk3 was non-functional (Fig. 3, A, D and E, and movie S3). Centrosome breakage was sudden and fast, with separation velocities in the range of micrometers per minute (Fig. 3, B and C), showing strong opposing forces that fracture the centrosome. Centrosome fracturing only occurred at path junctions, but not during migration along straight paths (Fig. 3, D and E, fig. S3A and movie S3). To confirm these results in complex but deformable 3D scaffolds, we analyzed the integrity of the centrosome during migration in collagen matrices. In the presence of GSK-626616, DCs showed broken pairs of centrioles, resulting in individual centrioles that even located in entirely different cellular parts (Fig. 3, F to H). Together, these findings discover that the centrosome can break mechanically and show that Dyrk3 maintains the mechanical integrity of the centrosome.

**Fig 3.**
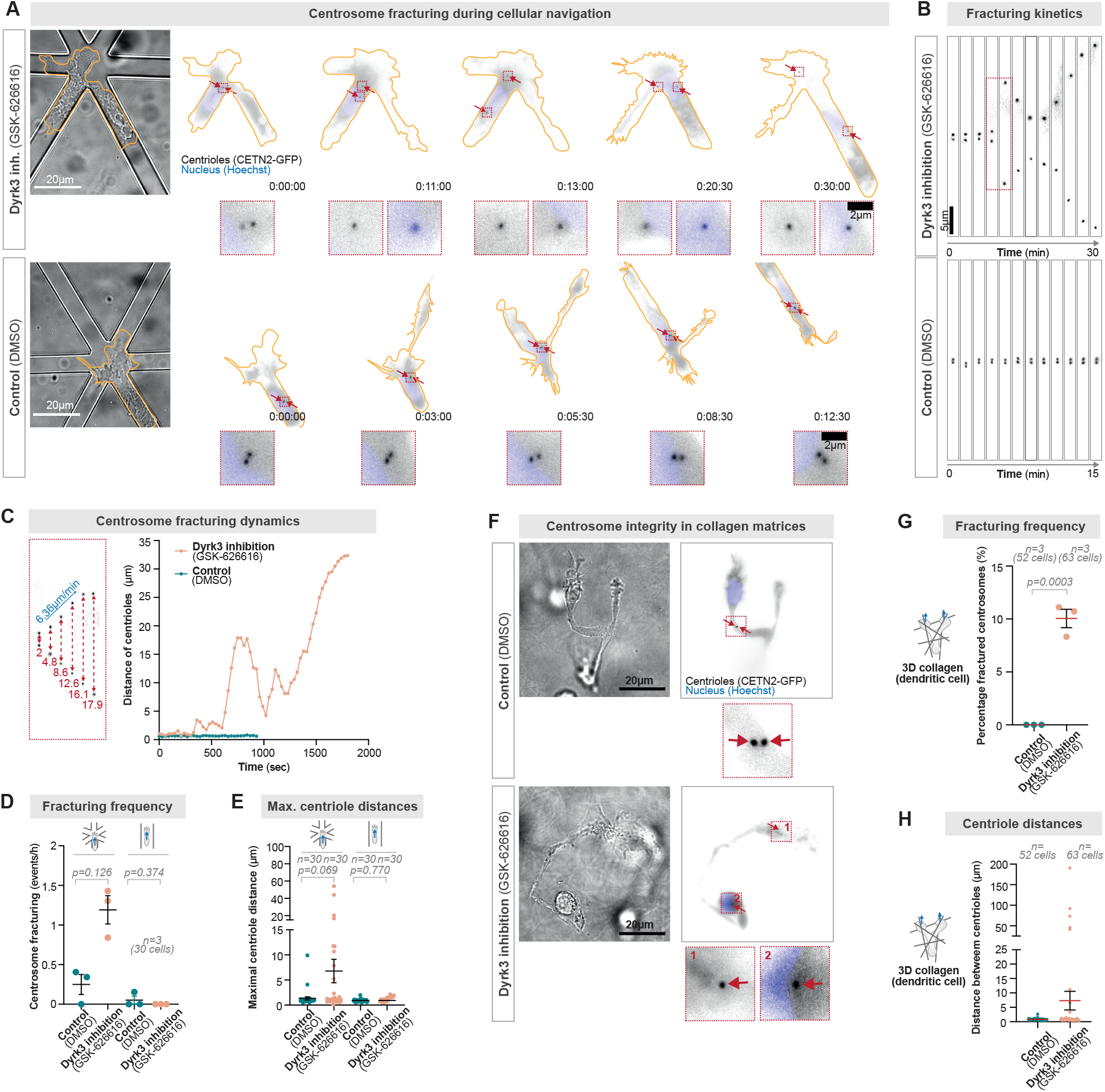
The centrosome fractures during cellular pathfinding. **(A)** Dynamics of the centriole pair (CETN2-GFP; black) during dendritic cell (DC) migration along a 6-way path junction in the presence of 5μM GSK-626616 or DMSO (control). Note the far-distant separation of the two centrioles in the presence of 5μM GSK-626616. The nucleus is stained with Hoechst (blue). Time is indicated as h:min:s. **(B)** As in (A), showing the detailed kinetics of the centriole pair. **(C)** As in (A), showing the detailed velocity of the separation of the centriole pair. **(D)** Frequency of centrosome breakage and **(E)** maximal distance of individual centrioles during DC migration along path junctions (6-way) or unidirectional paths. **(F)** Representative DC during migration in a deformable collagen matrix in the presence of 5μM GSK-626616 or DMSO (control). **(G)** Frequency of centrosome breakage and **(H)** maximal distance of individual centrioles during DC migration in 3D collagen matrices.

To disentangle extracellular from intracellular mechanical forces causing centrosome fracturing, we exposed the cells to strong environmental confinement by squeezing the cells between a layer of agarose and a coverslip (*21*). Centrosome fracturing occasionally occurred during this mechanical confinement in a Dyrk3-dependent manner (fig. S3, B and C). Similarly, we detected centrosome deformations during cellular squeezing through narrow microenvironmental pores, in particular during translocation through two-micrometer pores (fig. S4, A to D, and movie S4). Notably, when Dyrk3 was non-functional, centrosome stretching was already detectable during translocation through wider three- and four-micrometer pores (fig. S4, A to D, and movie S4). Thus, mechanical stresses from the microenvironment lead to centrosome deformations. Yet, as centrosome fracturing was more frequent during cellular path decisions, these findings suggested that intracellular forces from competing protrusions are an even larger source of mechanical centrosome deformations. Thus, we combined both forces at the same time, environmental confinement and competing cellular protrusions, by integrating small 6 micrometer-sized beads as path obstacles between the layer of agarose and the coverslip. In this confining and maze-like environment, CETN2-GFP expressing DCs showed high rates of centrosome fracturing when Dyrk3-was non-functional (fig. S3, D and E). Thus, intracellular as well as extracellular forces act together on the mechanical integrity of centrosomes.

Centrosome depletion often results in surprisingly mild cellular phenotypes, as other organelles such as Golgi membranes (*22*) are able to function as alternative microtubule organizing centers (MTOCs) (*4, 23*). Our findings of centrosome fracturing raised the possibility that the consequences of fractured centrosomes are entirely different to the consequences of centrosome non-functionality. To test this possibility, we visualized the microtubule cytoskeleton upon centrosome fracturing. Surprisingly, after fracturing of the centrosome, single distant centrioles often formed separate MTOCs, from which microtubules radially distributed in an aster-like manner (Fig. 4, A and B, and fig. S5A). In contrast, intact centriole pairs formed singular microtubule organizing centers that efficiently nucleated microtubules, even when Dyrk3 was inhibited (Fig. 4A, fig. S5, B to D, and movie S5). These results suggested the intriguing possibility that individual centrioles can form two coexisting functional MTOCs within one motile cell. To test their microtubule anchoring capacity, we imaged ninein as a major microtubule anchoring protein at centrioles (*7, 24*). In line with our observations that microtubules detached only occasionally (fig. S5E), ninein mostly localized to both individual centrioles upon centrosome fracturing (Fig. 4C, and fig. S6A), showing a functional microtubule anchoring capacity of both coexisting MTOCs. Similarly, the microtubule nucleator gamma-tubulin frequently localized around both individual centrioles after centrosome fracturing (Fig. 4D, and fig. S6B). Thus, fracturing of the centrosome leads to the emergence of two simultaneously coexisting functional microtubule organizing centers within a single cell.

**Fig 4.**
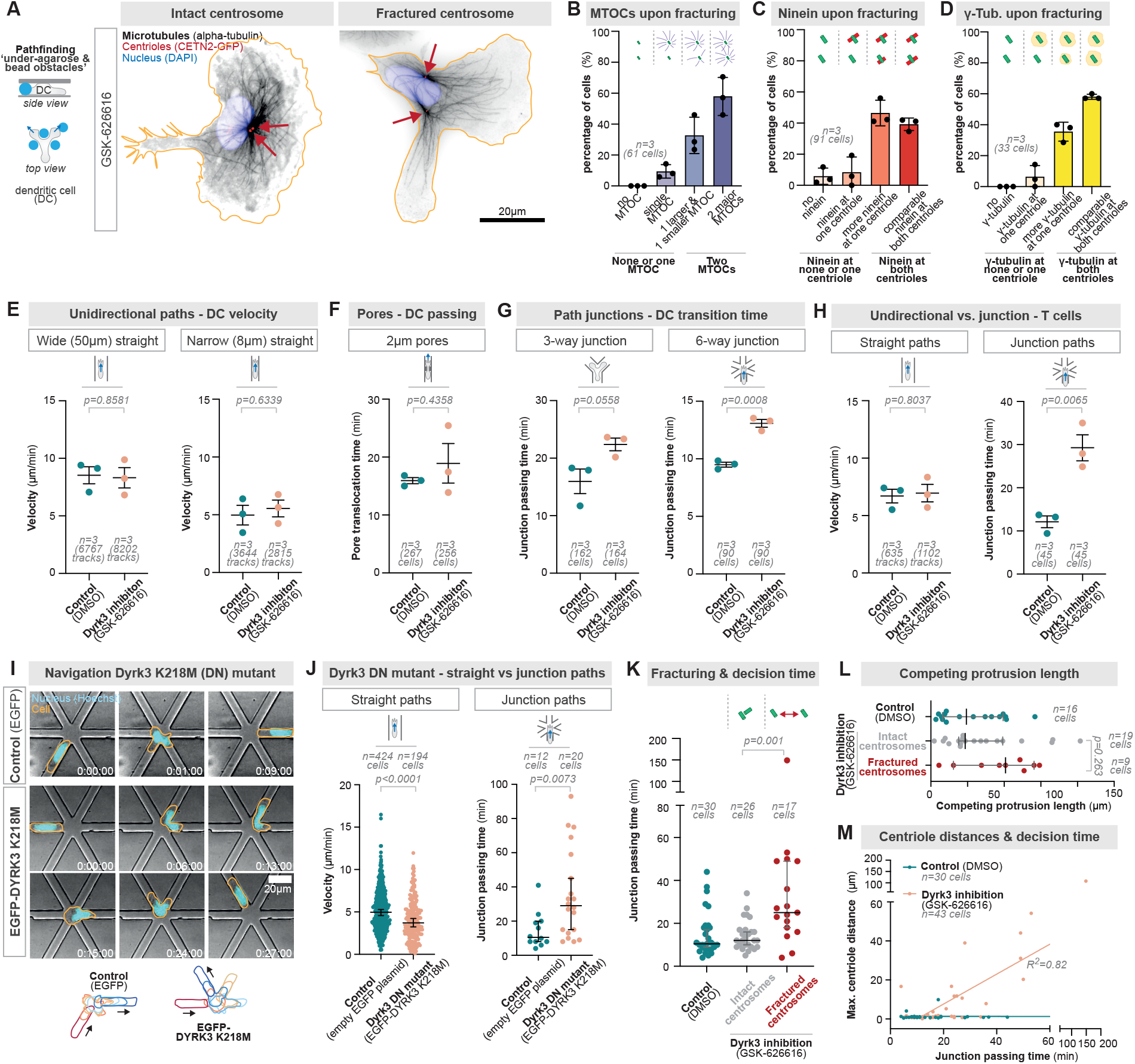
Centrosome fracturing impedes cellular navigation. **(A)** Immunofluorescence of Dyrk3-inhibited (5μM GSK-626616) CETN2-GFP (red, arrows) dendritic cells (DCs) upon migration through bead-obstacles underneath an agarose layer showing representative examples of intact and fractured centrosomes. Anti-alpha-tubulin (black) and DAPI (blue) visualize the microtubule cytoskeleton and the nucleus, respectively. **(B-D)** Quantification of microtubule aster formation (A), the microtubule anchoring protein ninein (B), and the microtubule nucleator gamma-tubulin (C) upon breakage of the centrosome. **(E-G)** DC velocity along unidirectional paths (E), 2 μm pores (F), and path junctions (G) in the presence of 5μM GSK-626616 or DMSO (control). **(H)** Jurkat T cell velocity along unidirectional paths or 6-way path junctions in the presence of 5μM GSK-626616 or DMSO (control). **(I)** Representative EGFP-Dyrk3 K218M or EGFP expressing Jurkat T cells migrating through a 6-way path junction. Time is indicated as h:min:s. **(J)** Quantification of I, in comparison to migration along straight paths. **(K)** Path decision time of CETN2-GFP DCs with fractured and non-fractured centrosomes through 6-way decision points. **(L)** Length of the major competing protrusion of DCs with fractured and non-fractured centrosomes migrating through 6-way junctions. **(M)** Correlation of the cellular decision time (as in K) with the maximal distance between individual centrioles.

Given the importance of singular microtubule organizing centers as a steering organelle during cellular locomotion (*11, 25*), we next tested the functional consequences of two coexisting MTOCs for moving cells. Rendering Dyrk3 non-functional by GSK-626616 and thus rendering the centrosome prone to fracturing, resulted in unaffected DC migration velocities along unidirectional straight paths that are wider or smaller (Fig. 4E, fig. S7, A and B, and movie S6) than the cell, as well as through narrow 2μm constrictions (Fig. 4F, fig. S7C, and movie S6). This is consistent with microtubules being dispensable for unidirectional cellular movement, as the actin cytoskeleton is the driving force for cellular forward locomotion (*8, 26*). In contrast, DCs moving through microchannels with either 3-way or 6-way path junctions needed longer to productively perform path decisions in the presence of GSK-626616 (Fig. 4G, fig. S7, D and E, and movie S6). To test the generality of these findings, we measured the migration properties of Jurkat T cells upon Dyrk3 inhibition with GSK-626616. Indeed, migration was only delayed at path junctions, while migration along unidirectional straight paths was unaffected (Fig. 4H, fig. S8, A and B, and movie S7). Similarly, expression of the kinase-dead point mutant of DYRK3 (K218M) only resulted in mildly reduced migration velocities along unidirectional straight paths, but it strongly delayed the passage through path junctions (Fig. 4, I and J, fig. S8, A and B, and movie S7). These data suggested that impaired centrosome stability hindered cell migration specifically during cellular navigation. Thus, we next measured CETN2-GFP DCs during path decisions, revealing that specifically cells with fractured centrosomes have longer competing protrusions (Fig. 4L) and require more time to productively make path decisions (Fig. 4K), where even the delay of passage correlated with the separation distance between fractured centrosome parts (Fig. 4M). Thus, mechanical centrosome fracturing impedes cell functionality by generating coexisting microtubule organizing centers that compete during cellular path navigation and thereby cause cellular entanglement in the microenvironmental matrix.

How membrane-less organelles (*27*) respond to mechanical forces and withstand them is largely unknown. The centrosome is a central cellular membrane-less organelle, which undergoes a highly regulated and timed separation during cell division. While the lack of a surrounding membrane is critical for this intended centriolar separation process at each and every cell division, our work discovers that centrosomes are subject to mechanical deformations in non-dividing cells by extracellular and intracellular forces, which are even able to disrupt centrosome integrity. To ensure and maintain mechanical stability, centrosomes likely evolved concurrent and synchronous intra-centriolar cohesion mechanisms, including a cNAP1- and rootletin-based protein linker directly inter-connecting the centriole pair, a microtubule-based connector that may even provide long-range connection when required, and presumably a PCM-based stabilizer surrounding the centriole pair (*12, 28*–*30*). Beyond its relevance for cells moving in multicellular organisms, such as during immune surveillance and cancer cell motility, our findings suggest the fundamental concept that cells may have to preserve the integrity of the centrosome and other membrane-less organelles whenever they experience mechanical forces, such as during development, tissue homeostasis, or disease.

## Acknowledgments

We thank Lukas Pelkmans and Dorothee Dormann for providing Dyrk3-EGFP plasmids, Michelle Duggan for RNA isolation from migrating DCs, Michael Schuster from the Biomedical Sequencing Facility at CeMM, Jan Schwarz for providing Jurkat T cells, Maximillian Götz for initial transcriptome analysis, Michael Sixt for long-standing mentoring and support, Malte Benjamin Braun for critical reading of the manuscript, and the Core Facility Bioimaging, the Core Facility Flow Cytometry, and the Animal Core Facility of the Biomedical Center (BMC) for excellent support.

## Funding

Peter Hans Hofschneider Professorship of the Stiftung Experimentelle Biomedizin (JR)

German Research Foundation grant ‘CRC914, project A12’ (JR)

German Research Foundation grant ‘SPP2332, project 492014049’ (JR)

LMU Institutional Strategy LMU-Excellent within the framework of the German Excellence Initiative (JR)

Deutsche Forschungsgemeinschaft (DFG, German Research Foundation) under Germany’s Excellence Strategy – EXC 2151 – 390873048 (EK)

## Author contributions

Conceptualization: MTS, JR

Methodology: MTS, MRF, JK, JM, EK

Image analysis software: RH

Transcriptomics & analysis: TP, CB, TS

Investigation: MTS, KS, JR

Funding acquisition: JR

Project administration: JR

Supervision: JR

Writing – original draft: MTS, JR

Writing – review & editing: MTS, MRF, JK, RH, JM, TP, EK, CB, TS, JR

## Data and materials availability

RNAseq data are available at the GEO repository (GSE220623). All other data are available in the main text or the supplementary materials.

