## Supplementary Materials for "Mechanical Centrosome Fracturing during Cell Navigation"

**This PDF file includes:**

Materials and Methods
Figs. S1 to S8
Captions for Movies S1 to S7

**Other Supplementary Materials for this manuscript include the following:**

Movies S1 to S7

#### Materials and Methods

Cell culture

All cells were grown and maintained at 37 °C in a humidified incubator with 5 % $CO_2$. Jurkat T cells were cultured in R10 medium at a cell density of 0.1–1.5x$10^6$ cells/ml. DCs were differentiated either from bone marrow isolated from male C57Bl6/J wildtype mice (aged 8-12 weeks) or from Hoxb8 (*31*) precursor cell lines (CETN2-GFP (*10*) and EB3-mCherry (*11*) ). Cells were cultured in R10 medium (RPMI1640 + 10 % fetal calf serum (FCS), 2 mM L-glutamine, 100 U/ml penicillin, 100 mg/ml streptomycin and 0.1 mM 2-mercaptoethanol, all Gibco) supplemented with 10 % granulocyte–macrophage colony-stimulating factor (GM-CSF) hybridoma supernatant. On differentiation days 3 and 6, fresh medium was added. For migration experiments, either fresh or thawed DCs (differentiation day 8) were stimulated by adding 200 ng/ml lipopolysaccharide (LPS; *E. coli* O26:B6, MilliporeSigma) for 24 h to induce cell maturation.

Mice

All animals were housed in the Core Facility Animal Models at the Biomedical Centre (Ludwig-Maximilians-Universität) and animal procedures and experiments were in accordance with the ministry of animal welfare of the region of Oberbayern and with the German law of animal welfare.

Flow cytometry analysis

DC maturation was routinely checked for surface expression of CD11c and MHCII. After Fc receptor blockage using an anti-mouse CD16/32 antibody (14-0161-85, Invitrogen) diluted in FACS buffer (1 % BSA, 2 mM EDTA in PBS), cells were stained with anti-mouse CD11c and anti-mouse MHCII antibodies (17-0114-82, 48-5321-82, both Invitrogen). Flow cytometry analysis was performed on a Cytoflex S flow cytometer (Beckmann-Coulter).

Transgene delivery

For transient expression of fluorescent reporter constructs, Jurkat T cells were electroporated with plasmids encoding the respective constructs 16 hours prior to the experiment using the Neon Transfection system (Invitrogen) with 3 pulses at 1600 V for 10 ms each.

Dyrk3 localization was analyzed by co-electroporating Jurkat T cells with pcDNA5-Dyrk3-WT-GFP (*20*) (a gift from Lucas Pelkmans and Dorothee Dormann), and mCherry-Centrin2-N-10 (a gift from Michael Davidson; Addgene plasmid # 55018). The next day, cells were injected in an under-agarose migration assay as described below and mCherry$^+$ GFP$^+$ cells were imaged.

For validation of Dyrk3 inhibition mediated effects on migration, Jurkat T cells were transfected either with pcDNA5/ FRT/ TO- GFP (*32*) (a gift from Harm Kampinga; Addgene plasmid # 19444) as control, or pcDNA5-Dyrk3-K218M-GFP (*20*) (a gift from Lucas Pelkmans and Dorothee Dormann) encoding for a dominant-negative point-mutant Dyrk3, respectively. The next day, cells were prepared for fluorescence-activated cell sorting on a FACSAriaFusion (BD) equipped with 4 lasers (405, 488, 561, 640 nm). Live GFP$^+$ cells were directly sorted into R10 medium buffered with 25 mM HEPES. After 1 hour of recovery in the incubator, cells were used for downstream migration assays as described below.

Micro-fabricated devices

Micro-fabricated devices were prepared as described previously (*33*). Briefly, micro-structures were replicated from custom-made wafers produced by photolithography or epoxy replicates, with defined width, height, length, and pore sizes. The height of the micro-structures ranged between 4-5 µm to allow cell confinement from top and bottom. Wide straight channels had a width of 50 µm. Narrow straight channels, channels with constrictions, 3-way pathfinding channels, and 6-way pathfinding channels had a width of 8 µm, thereby confining cells from all sides. Distance between two 6-way crossings was 90 µm. The pore size of microchannels with constrictions was 2, 3, or 4 µm, as indicated.

Polydimethylsiloxane (PDMS, 10:1 mixture of Sylgard 184, Biesterfeld) was added onto the template structures to generate replica of the micro-structures. Air bubbles were removed with a desiccator. After solidification at 80 °C overnight, PDMS was carefully removed from the templates and cut into pieces according to the respective design size. Holes for cell and chemokine loading were punched. Using a plasma cleaner, the PDMS device was bonded to clean glass coverslips. PDMS devices were then placed at 120 °C for 10 min, followed by overnight incubation at 80 °C to permanently bond them to the glass surface.

Live-Cell migration assays

For live cell imaging, cell nuclei were visualized by pre-incubating cells for at least 30 min with NucBlue (Invitrogen) according to the manufacturer's instructions, followed by washing. For pharmacological inhibition experiments, final concentrations of 1-10 µM GSK-616626 (Tocris; dissolved in DMSO) and 10-100 µM Harmine (Sigma; dissolved in DMSO) were used as indicated. Control samples were treated with DMSO in the corresponding dilution.

*Micro-channel migration assays:* Prior to the experiment, PDMS devices were flushed with phenol-free R10 medium supplemented with 50 µM L-ascorbic acid (MilliporeSigma) and inhibitors if needed according to the experimental set-up. After incubation at 37 °C, 5 % $CO_2$ in a cell culture incubator for at least 1 hour, devices were used for downstream experiments. Then, 0.625 µg/ml CCL19 (DCs) or 1.25 µg/ml CXCL12 (T cells) were loaded into the chemokine loading hole to establish a chemokine gradient, followed by addition of 0.3-0.5x$10^5$ cells into the second loading hole.

*Under-agarose migration assays:* Under-agarose migration assays without bead obstacles were prepared as described previously (*34*). Briefly, 1 % agarose was prepared by mixing 4 % UltraPure agarose (Invitrogen) in sterile water with 55 °C prewarmed phenol-free RPMI-1640 (Gibco) supplemented with 20 % FCS, 1x Hanks buffered salt solution pH 7.3 in a 1:3 ratio. For experiments including inhibitors, the 1 % agarose mixture was let cool down to 37 °C before adding the inhibitor to the respective final concentration. The agarose was poured into imaging-suitable 8-well slides (Ibidi), polymerized for 1 hour at room temperature and was then transferred to the incubator for 1 hour for equilibration. For under-agarose migration assays including bead obstacles, 8-well slides were pre-coated with Polybeads® (6 µm diameter, 07312-5, Polysciences). The Polybeads® were washed and resuspended in phenol-free R10 medium. After activating the well glass surface with oxygen plasma, beads were added for coating. Then, the agarose mixture was added after additional 30 min to ensure stable bead attachment. For both, under agarose with and without bead obstacles, 2 mm wide holes were generated using tissue biopsy punchers after agarose solidification and equilibration. Then, CCL19 (2.5 µg/ ml; DCs) or CXCL12 (5 µg/ml; T cells) in phenol-free R10 were loaded as chemotactic stimulus. 0.2x$10^5$ cells were injected between the glass surface and agarose layer in a distance of 2-3 mm to the chemokine loading hole. For live-imaging, assays were placed in the incubator for 1 hour to allow induction of directional migration towards the chemokine source before imaging.

*Collagen migration assays:* collagen migration assays were performed as described previously (*33*, *35*). Briefly, for DC collagen migration assays, PureCol bovine collagen (Advanced BioMatrix) in 1× minimum essential medium (MEM, Sigma) and 0.4 % sodium bicarbonate (Sigma) was mixed with $3x10^5$ cells in R10 at a 2:1 ratio, resulting in gels with a collagen concentration of 1.7 mg/ml. Collagen-cell mixtures were cast in custom-made migration chambers with a diameter of 18 mm and a height of approx. 1 mm. After polymerization of collagen fibers at 37 °C, 5 % $CO_2$ in a cell culture incubator for 75 min, 80 µl CCL19 (0.625 µg/ml, 440-M3-025, Bio-Techne) was added to the top of the chamber. For Jurkat T cell collagen migration assays, PureCol stock solution was diluted with PBS, resulting in a final collagen concentration of 1.3 mg/ml mixed with $2x10^5$ cells in R10 at a 2:1 ratio. After polymerization at 37 °C, 5 % $CO_2$ in a cell culture incubator for 75 min, 80 µl CXCL12 (1.25 µg/ml, 350-NS-050, Bio-Techne) was added to the top of the chamber. For inhibition experiments, inhibitors were added to the collagen-cell mixture as well as to the chemokine solution at the indicated concentrations.

Immunofluorescence stainings

For immunofluorescence stainings, under-agarose migration assays were prepared as described above. Following cell migration for 2 hours, 3.7 % paraformaldehyde (PFA; diluted in PBS) prewarmed to 37 °C was added on top of the agarose and incubated for 1 hour at 37 °C, 5 % $CO_2$. After fixation the agarose block was carefully removed and cells were washed with PBS. Following permeabilization with 1x SAPO buffer (0.2% BSA + 0.05% saponin diluted in PBS) for 30 min, blocking was performed with 5 % BSA (diluted in 1x SAPO). Primary antibodies were incubated overnight at 4 °C (rat anti-α-tubulin: 2 µg/ml, MA1-80017, Invitrogen; rabbit anti-ninein: 0.25 µg/ml, PA5-82224, Invitrogen; mouse anti-γ-tubulin: 4.25 µg/ml, T6557, Sigma; all diluted in 1x SAPO). The next day, samples were washed with PBS and stained with secondary antibodies (goat anti-rat Alexa Fluor® Plus 647: 4 µg/ml, A48265, Invitrogen; donkey anti-rabbit Alexa Fluor® Plus 647: 4 µg/ml, A32795, Invitrogen, goat anti-mouse Alexa Fluor® 555: 2 µg/ml, ab150114, abcam; all diluted in 1x SAPO) and DAPI (1:10000, Thermo Fisher Scientific) at room temperature for 1 hour. After washing with PBS, cells were mounted using Fluoromount-G (Invitrogen).

Transcriptomics of migrating dendritic cells

Bone-marrow derived dendritic cells were incorporated into three-dimensional collagen gels as described above. Collagen gels of different densities were obtained by using different bovine collagen stock dilutions to final concentrations of 1.7 mg/ml, 2.6 mg/ml, 3.5 mg/ml, and 4.4 mg/ml. Controls were performed by placing DCs into the same migration chambers but without collagen. Five hours after introduction of the CCL19 chemokine gradient, the gel was isolated from the chamber and immediately bathed in Trizol. Subsequently, RNA extraction was performed following the 'Total RNA extraction' protocol of the 'Immunological Genome Project' (ImmGen.org). The amount of total RNA was quantified using the Qubit 2.0 Fluorometric Quantitation system (Thermo Fisher Scientific, Waltham, MA, USA) and the RNA integrity number (RIN) was determined using the Experion Automated Electrophoresis System (Bio-Rad, Hercules, CA, USA). RNA-seq libraries were prepared with the TruSeq Stranded mRNA LT sample preparation kit (Illumina, San Diego, CA, USA) using Sciclone and Zephyr liquid handling workstations (PerkinElmer, Waltham, MA, USA) for pre- and post-PCR steps, respectively. Library concentrations were quantified with the Qubit 2.0 Fluorometric Quantitation system (Life Technologies, Carlsbad, CA, USA) and the size distribution was assessed using the Experion Automated Electrophoresis System (Bio-Rad, Hercules, CA, USA). Before sequencing on an Illumina HiSeq 2000 instrument following a 50 base single end protocol, samples were diluted and pooled into NGS libraries in equimolar amounts. For analysis, sequencing reads were aligned to the mouse reference genome (version GRCm38.95) with STAR (version 2.7.0f). Expression values (TPM) were calculated with RSEM (version 1.3.0). Post-processing was performed in R/bioconductor (version 3.5.3) using default parameters if not indicated otherwise. Differential gene expression analysis was performed with 'DEseq2' (version 1.22.2). An adjusted p value (FDR) of less than 0.1 was used to classify significantly changed expression.

Imaging

Live-cell imaging was performed at 37 °C, and supplementation with 5 % $CO_2$ in a humidified chamber if needed. Cell migration was recorded using conventional inverted wide-field DMi8 microscopes (Leica) using HC PL FLUOTAR 4×/0.5 PH0 air, HC PL FLUOTAR L20x/0.40 PH1 air, HC PL APO 40×/0.9 PH3 air and HC PL APO 100x/1.40 oil objectives, equipped with a Lumencor or pE-4000 light source (395, 475, 555, and 635 nm) and an incubation chamber, heated stage and $CO_2$ mixer (Pecon). For evaluation of microtubule nucleation dynamics, EB3-mCherry expressing DCs were imaged at 5 s intervals. Acquisition of immunofluorescence samples was performed on an inverted wide-field DMi8 microscopes (Leica) equipped with an HC PL APO 100x/1.47 oil objective.

Image analysis

Fiji / ImageJ (*36*) and Imaris (Bitplane) were used for image processing. In general, only single, non-interacting cells were included for analysis to avoid effects of neighboring cells on cell path, cell speed and centriolar dynamics.

Velocity of migrating DCs and Jurkat T cells along unidirectional paths (narrow straight, wide straight micro-channels) was analyzed using the tracking function of Imaris v9.7.2. For DC migration, cell nuclei were tracked with the following settings: object diameter 12 μm, manually adjusted quality threshold, autoregressive motion tracking algorithm (max. distance 25 μm, gap size =3), min. track duration 15 min. For Jurkat T cell migration the following settings were applied: object diameter 15 μm, manually adjusted quality threshold, autoregressive motion tracking algorithm (max. distance 15 μm, gap size=3), min. track duration 15 min. Pore translocation time and junction passing time, as well as centriolar distances were manually quantified in Fiji.

Centrosome splitting (fracturing) was classified as a distance above 1.5 μm between the two individual centrioles in a centriolar pair. For characterization of fractured centrosomes by immunostainings, recorded cells were analyzed by categorizing fluorescence signal patterns as indicated. Dyrk3 localization was evaluated by measuring fluorescence intensity profiles along a 5 μm long line determined by the centriolar axis using the Plot profile function in Fiji. Then, fluorescence values were normalized to the mean intensity value of the first and last four measured values, respectively. The competing protrusion length was measured using ImageJ by determining the maximal length for each protrusion during a productive path decision. The longest of those protrusions was defined as the main competing protrusion.

For analysis of microtubule nucleation rate and speed, cells that were well separated from other cells were randomly selected and cut out from raw movies. The EB3 comets within these cells were then tracked in Fiji using TrackMate v7.9.2 (*37*) (https://doi.org/10.1038/s41592-022-01507-1) with the following settings: LoG detector (object diameter: 0.65 µm), manually adjusted quality threshold and min. intensity filters, Kalman tracker: search radii 7,10 µm , no frame gap). The resulting tracks were exported and the presented statistics were derived with a custom Matlab script.

DC migration in collagen matrices was analyzed using a custom-made cell tracking tool for ImageJ (*14*). In brief, cell migration image sequences were background corrected by subtracting the average of the entire sequence. Particle filtering was used to discard objects smaller or larger than the cells. Then, for each image in the sequence the lateral displacement that optimizes its overlap with the previous frame was determined. Finally, the migration velocity towards the chemokine source was calculated from the y-displacement and the time between two consecutive frames. To analyze Jurkat T cell migration, cells were manually tracked in Fiji / ImageJ using the manual tracking plugin. Migration speed, accumulated distance and directionality were calculated using the Ibidi chemotaxis and migration tool (*38*). The first 30 minutes of the recordings were excluded from analysis due to initial image drift.

Statistics

All data that show individual cellular data points derive from cells from at least three independent biological replicates. All replicates were validated independently and pooled only when all showed similar results. Statistical analysis was conducted using GraphPad Prism using the appropriate tests according to normal or non-normal data distribution: unpaired t-tests (Fig. 3, D and G, Fig. 4, E to H, and fig. S3, B to E), Mann-Whitney (Fig. 2, C and D, Fig. 3E, and Fig. 4, J to L), Kruskal-Wallis with Dunn's multiple comparison test (Fig. 1, E and F), and linear regression fit (Fig. 4L). Error bars represent mean +/- SD (Fig. 4, B to D), mean +/- SEM (Fig. 2, A and B, Fig. 3, D, E, G, H, and Fig. 4, E to H), mean +/- 95% CI (Fig. 2G, and fig. S3, B to E), and median +/- 95% CI (Fig. 2, C and D, Fig. 4, J to L, and fig. S2D).

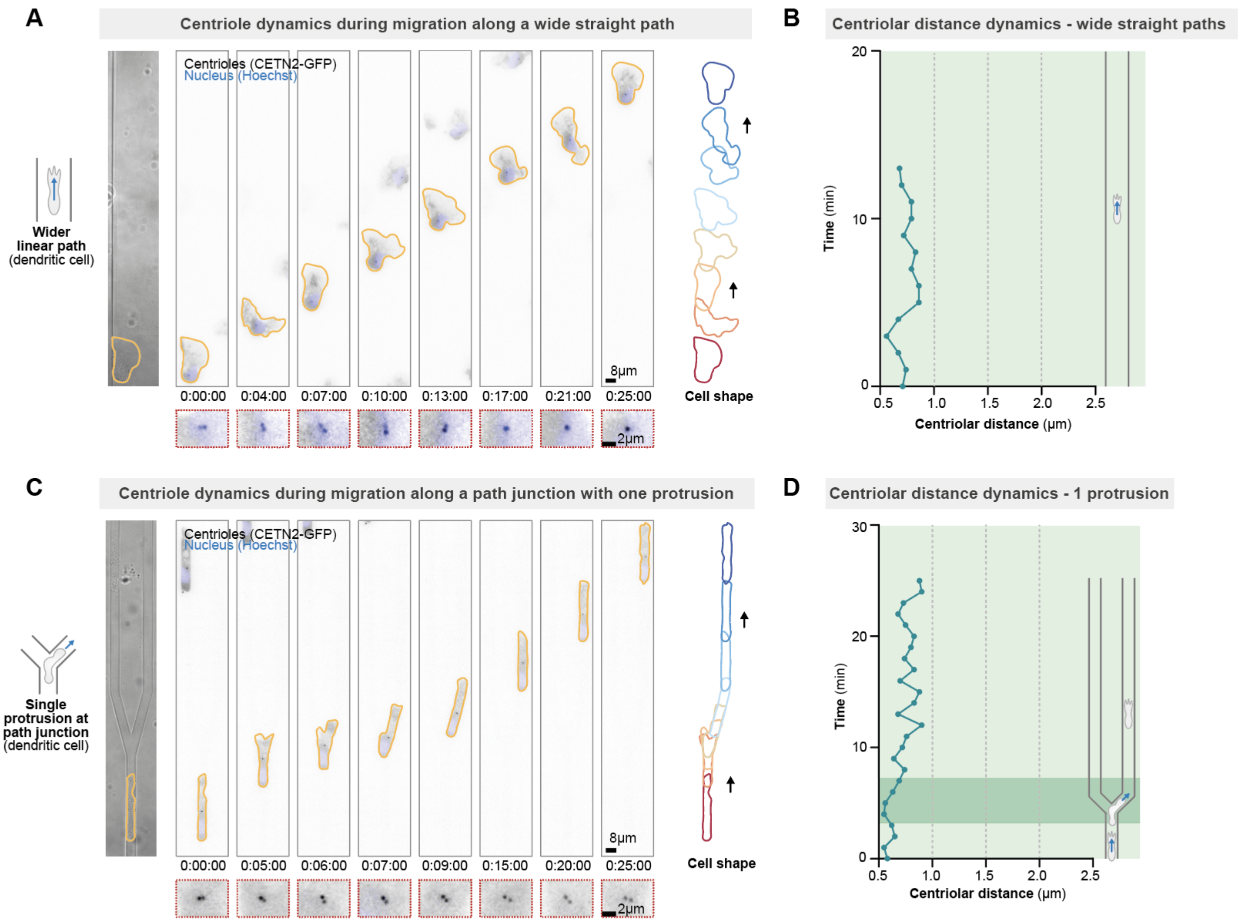


**Fig. S1.**

Dynamics of the centriole pair during DC migration along unidirectional paths. **(A)** Representative CETN2-GFP (centriole pair; black; enlargement in red dashed boxes) expressing dendritic cell (DC) stained with Hoechst (nucleus; blue) migrating along a unidirectional straight path (50 µm wide linear microchannel). **(B)** Centriolar distance dynamics during migration as depicted in (A); note the stable proximity of the individual centrioles. See Fig. 1E for quantification of centriolar distances during migration in this microenvironment composed of wide straight paths. **(C)** Representative CETN2-GFP (centriole pair; black; enlargement in red dashed boxes) expressing DC stained with Hoechst (nucleus; blue) migrating through a path junction (Y microchannel). Note that this specific cell has only one cell front at the path junction, and directly follows this cell protrusion into one of the two alterative paths. See Fig. 1F for quantification of centriolar distances while cells explore the alternative paths with one protrusion (and in comparison, to two protrusions). All data show representative cells from at least three independent biological replicates. Time is indicated as h:min:s.

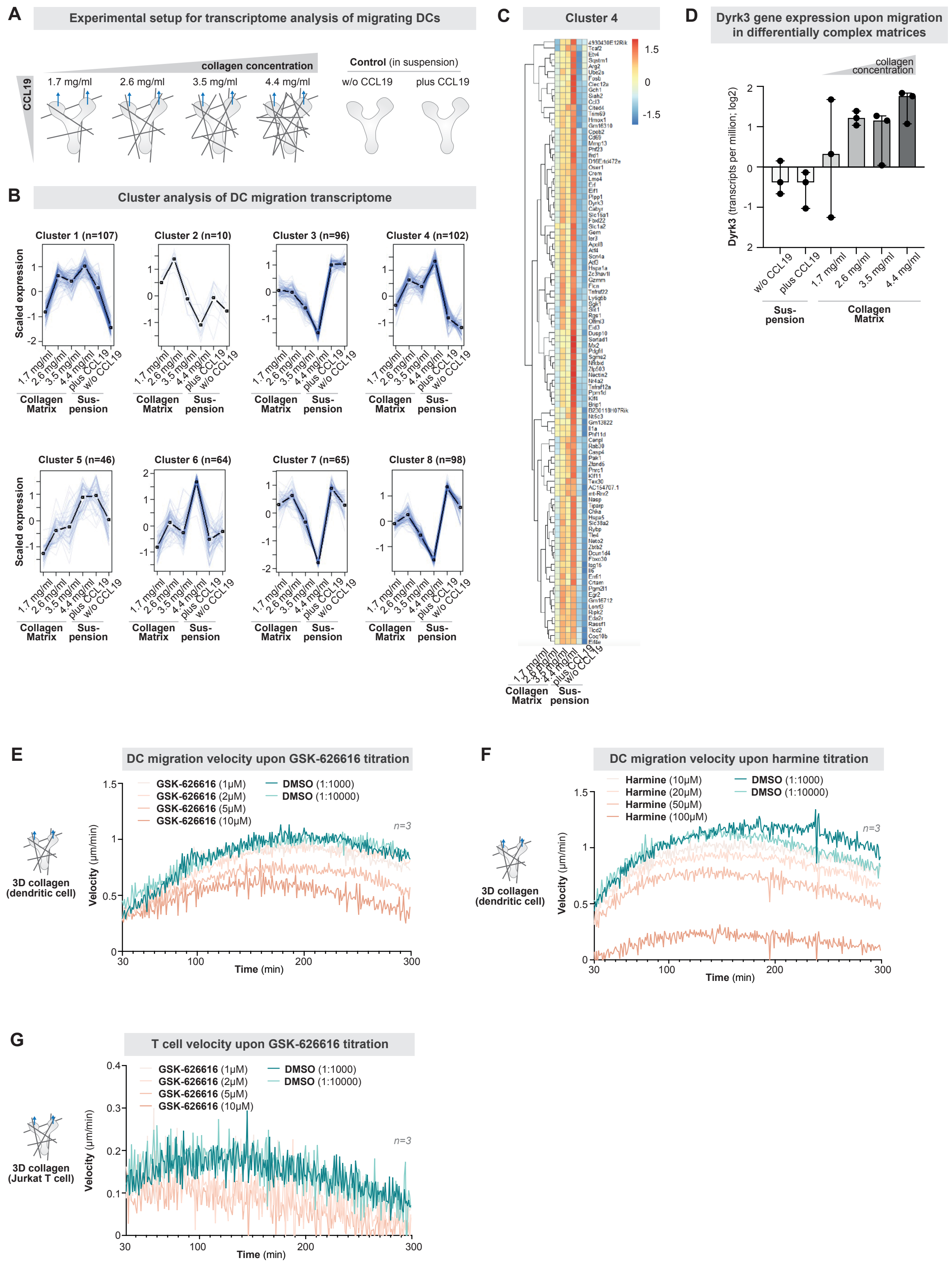


**Fig. S2.**

Transcriptomics of migrating dendritic cells and pharmacological inhibition of Dyrk3 during dendritic cell and T cell migration. **(A)** Scheme of the experimental setup to investigate the transcriptome of migrating dendritic cells in differentially complex collagen matrices (see 'Materials and Methods' for details). **(B)** Cluster analysis of differentially expressed genes derived from (A). **(C)** Close up view of genes in cluster 4, which upregulate their gene expression in more complex collagen matrices. **(D)** Differential gene expression of Dyrk3 (n=3 experiments; median; 95% confidence interval). **(E)** Dendritic cell migration in three-dimensional (3D) collagen matrices (1.7 mg/ml) along a CCL19 chemokine gradient in the presence of different concentrations of GSK-626616 (Dyrk3 inhibitor) or DMSO (control). The dataset for 5μM GSK-626616 is also shown in main Figure 2A. **(F)** Dendritic cell migration in three-dimensional (3D) collagen matrices (1.7 mg/ml) along a CCL19 chemokine gradient in the presence of different concentrations of Harmine (Dyrk inhibitor) or DMSO (control). The dataset for 50μM Harmine is also shown in main Figure 2B. **(G)** Jurkat T cell migration in three-dimensional (3D) collagen matrices (1.3 mg/ml) along a CXCL12 chemokine gradient in the presence of different concentrations of GSK-626616 (Dyrk3 inhibitor) or DMSO (control). All data derive from at least three independent biological replicates.

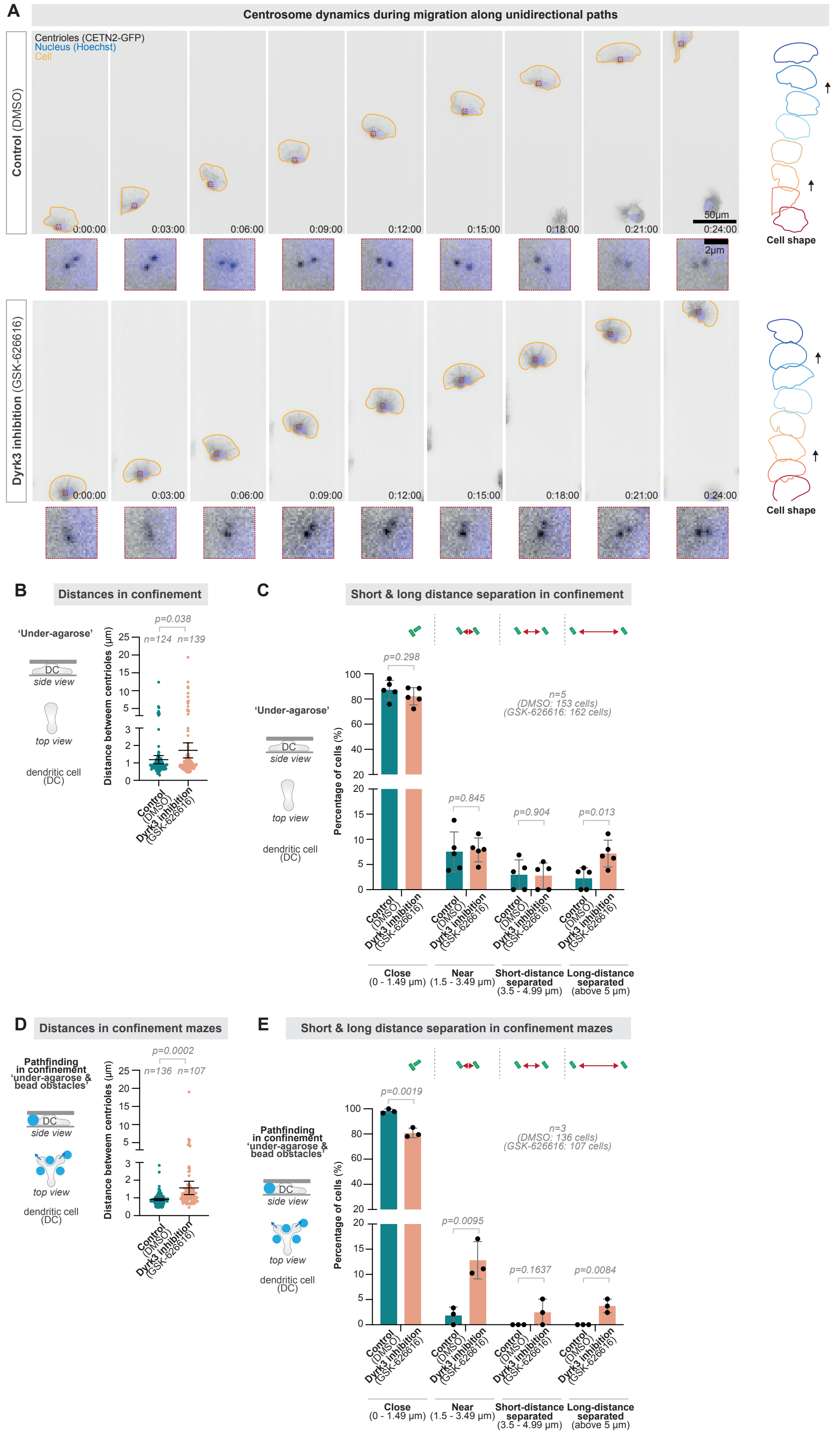

A
Centrosome dynamics during migration along unidirectional paths
Centrioles (CETN2-GFP)
Nucleus (Hoechst)
Cell
Control (DMSO)
Dyrk3 inhibition (GSK-626616)
0:00:00
0:03:00
0:06:00
0:09:00
0:12:00
0:15:00
0:18:00
0:21:00
0:24:00
50µm
2µm
Cell shape
B
Distances in confinement
'Under-agarose'
DC
side view
top view
dendritic cell (DC)
p=0.038
n=124
n=139
Distance betweem centrioles (µm)
Control (DMSO)
Dyrk3 inhibition (GSK-626616)
C
Short & long distance separation in confinement
p=0.298
p=0.845
p=0.904
p=0.013
n=5 (DMSO: 153 cells) (GSK-626616: 162 cells)
Percentage of cells (%)
Close (0 - 1.49 µm)
Near (1.5 - 3.49 µm)
Short-distance separated (3.5 - 4.99 µm)
Long-distance separated (above 5 µm)
D
Distances in confinement mazes
Pathfinding in confinement 'under-agarose & bead obstacles'
p=0.0002
n=136
n=107
E
Short & long distance separation in confinement mazes
p=0.0019
p=0.0095
p=0.1637
p=0.0084
n=3 (DMSO: 136 cells) (GSK-626616: 107 cells)


### Fig. S3.

Centrosome deformation dynamics and fracturing frequency during migration along unidirectional paths, in microenvironmental confinement, and in confining mazes. **(A)** Representative CETN2-GFP (centriole pair; black; enlargement in red dashed boxes) expressing dendritic cells (DCs) stained with Hoechst (nucleus; blue) migrating along a unidirectional straight path (linear microchannel) in the presence of 5μM GSK-626616 or DMSO (control). See Figures 3D and 3E for quantification. **(B)** CETN2-GFP expressing DCs stained with Hoechst migrating in microenvironmental confinement ('under-agarose assay'), quantifying the distances between individual centrioles in the centriole pair. **(C)** As in (B), classifying the short- and long-distantly separated individual centrioles. **(D)** CETN2-GFP expressing DCs stained with Hoechst migrating in confining and maze-like microenvironments ('under-agarose assay with bead-obstacles'). **(E)** As in (D), quantifying the distances between individual centrioles in the centriole pair. All data derive from at least three independent biological replicates. Time is indicated as h:min:s.

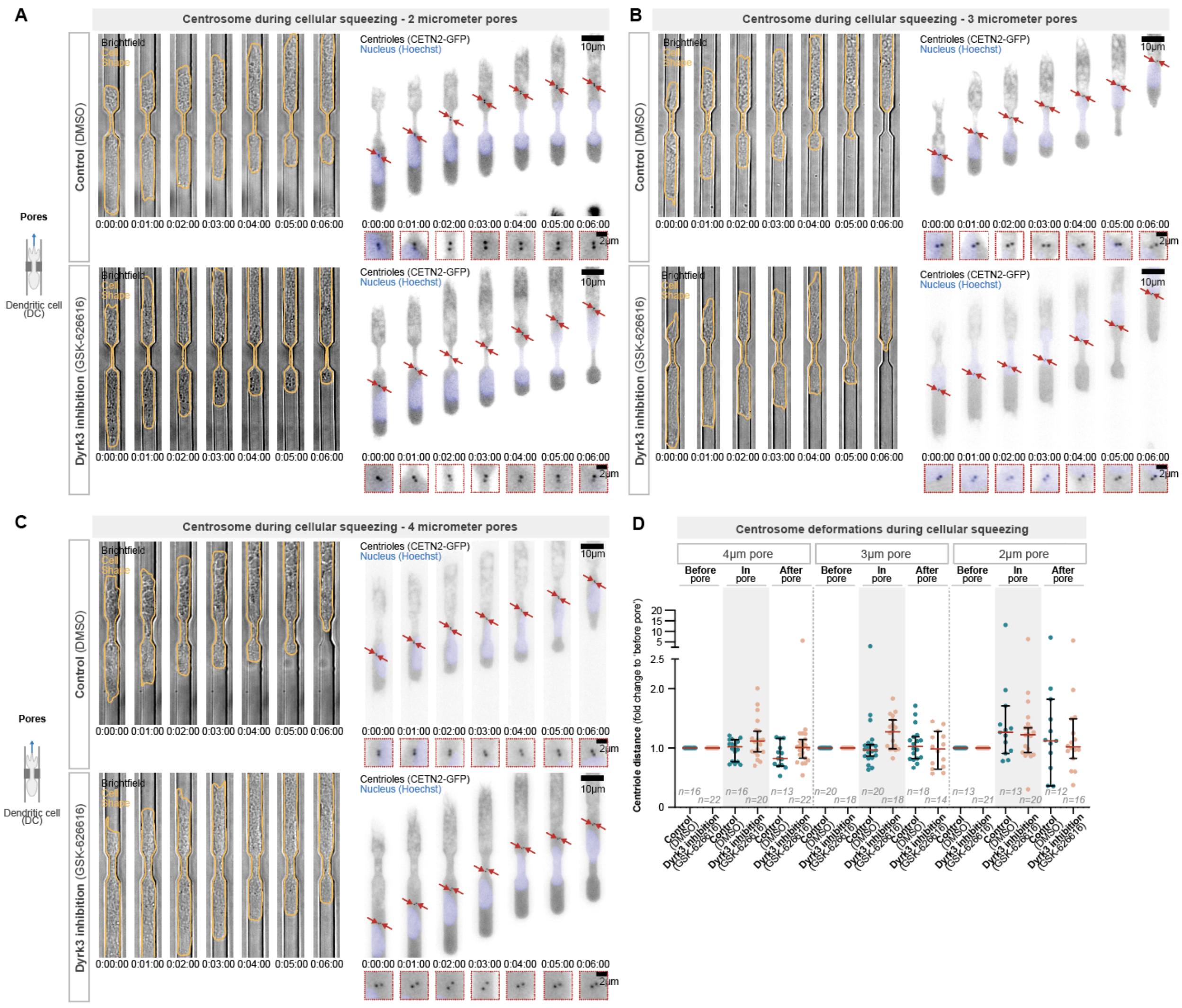


**Fig. S4.**

Centrosome deformation during cellular squeezing. **(A)** Representative CETN2-GFP (centriole pair; black; enlargement in red dashed boxes) expressing dendritic cells (DCs) stained with Hoechst (nucleus; blue) migrating along a single 2 µm pore (in a linear microchannel) in the presence of 5µM GSK-626616 or DMSO (control). **(B)** As in (A), but translocation through a 3 µm pore. **(C)** As in (A), but translocation through a 4 µm pore. **(D)** Quantification of centriolar distances in (A) to (C). All data show representative cells from at least three independent biological replicates. Time is indicated as h:min:s.

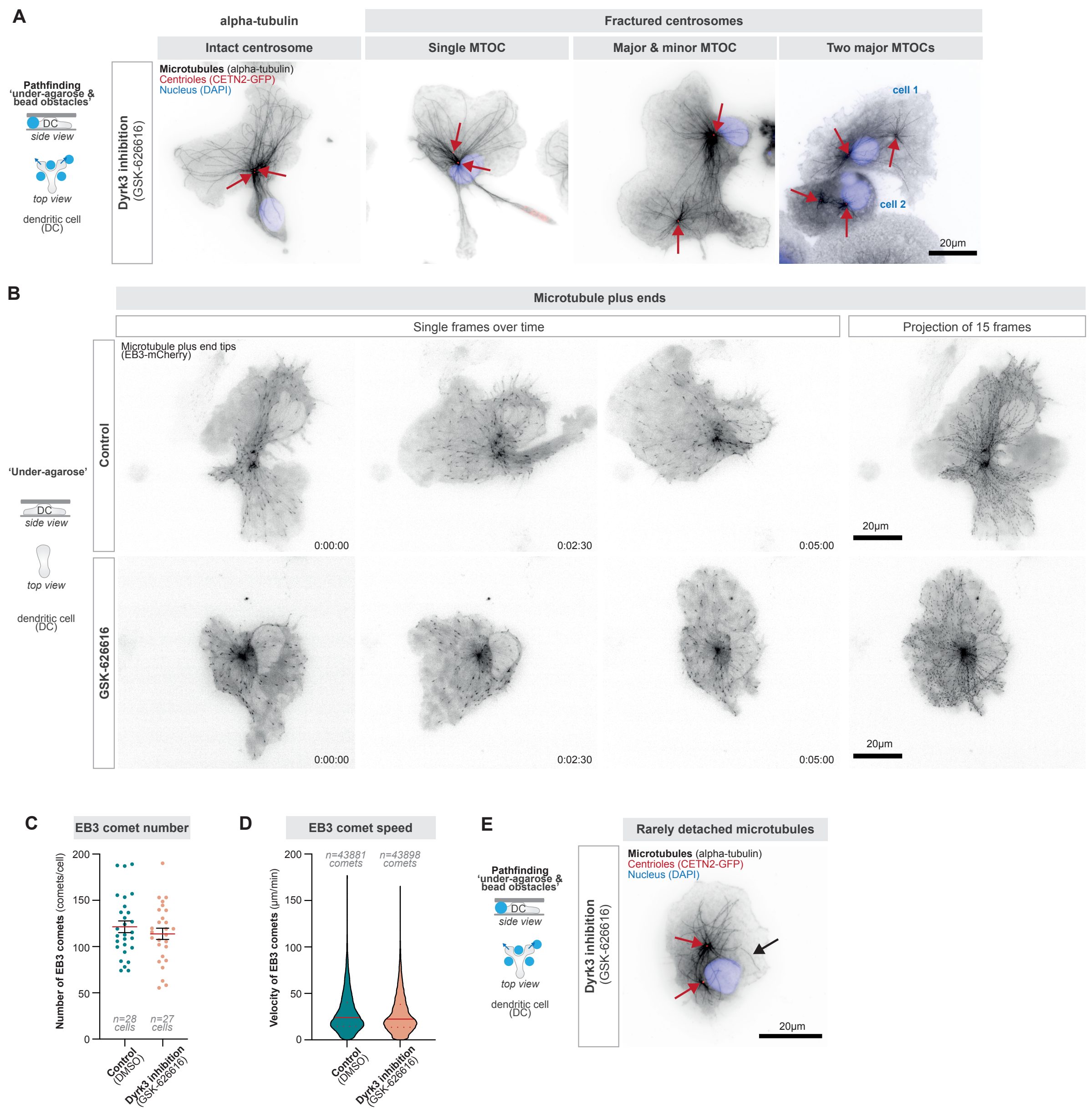


### Fig. S5.

The microtubule cytoskeleton upon Dyrk3 inhibition and centrosome fracturing. **(A)** Immunofluorescence staining of a representative CETN2-GFP (centriole pair; red) expressing dendritic cells (DCs) stained with DAPI (nucleus; blue) and with alpha-tubulin (black) in the presence of 5 µM GSK-626616. See Figure 4B for quantification. **(B)** Live-cell imaging examples of representative EB3-mCherry (microtubule-plus end tip marker; black) expressing DCs in the presence of 5 µM GSK-626616 or DMSO (control). Time is indicated as h:min:s. **(C)** Quantification of EB3 comets (microtubule plus ends) as shown in (B). **(D)** Quantification of EB3 comet velocity. **(E)** Exemplary DC with a rather rare event of a likely non-anchored microtubule (black arrow); CETN2-GFP (centriole pair; red) expressing DCs stained with DAPI (nucleus; blue) and with anti-alpha-tubulin (black) in the presence of 5 µM GSK-626616. All data show representative cells from at least three independent biological replicates.

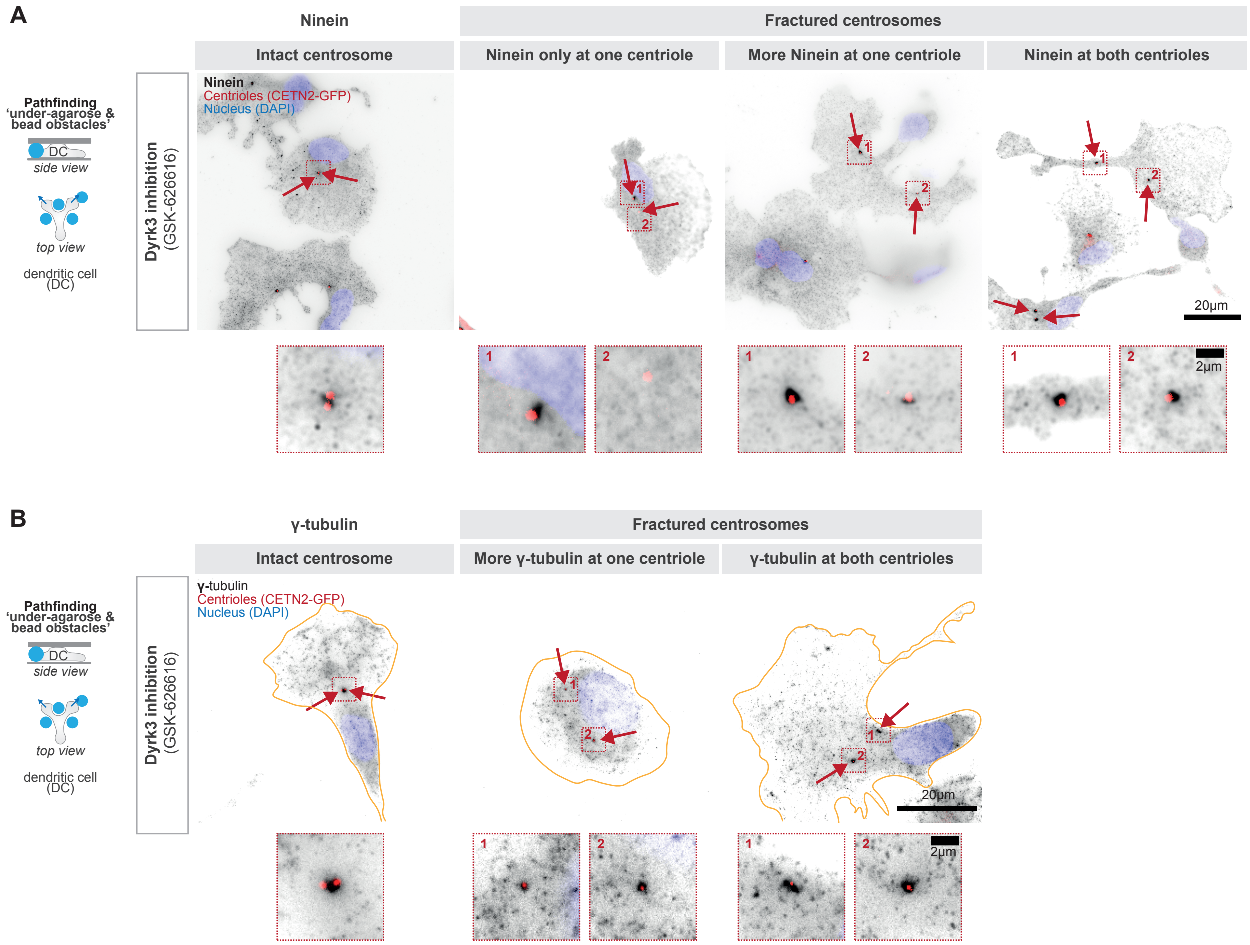


**Fig. S6.**

Microtubule anchoring and nucleating proteins upon Dyrk3 inhibition and upon centrosome fracturing. **(A)** Immunofluorescence staining of a representative CETN2-GFP (centriole pair; red) expressing dendritic cells (DCs) stained with DAPI (nucleus; blue) and with anti-ninein (black) in the presence of 5µM GSK-626616. See Figure 4C for quantification. **(B)** Immunofluorescence staining of a representative CETN2-GFP (centriole pair; red) expressing dendritic cells (DCs) stained with DAPI (nucleus; blue) and with anti-gamma-tubulin (black) in the presence of 5µM GSK-626616. See Figure 4D for quantification. All data show representative cells from at least three independent biological replicates.

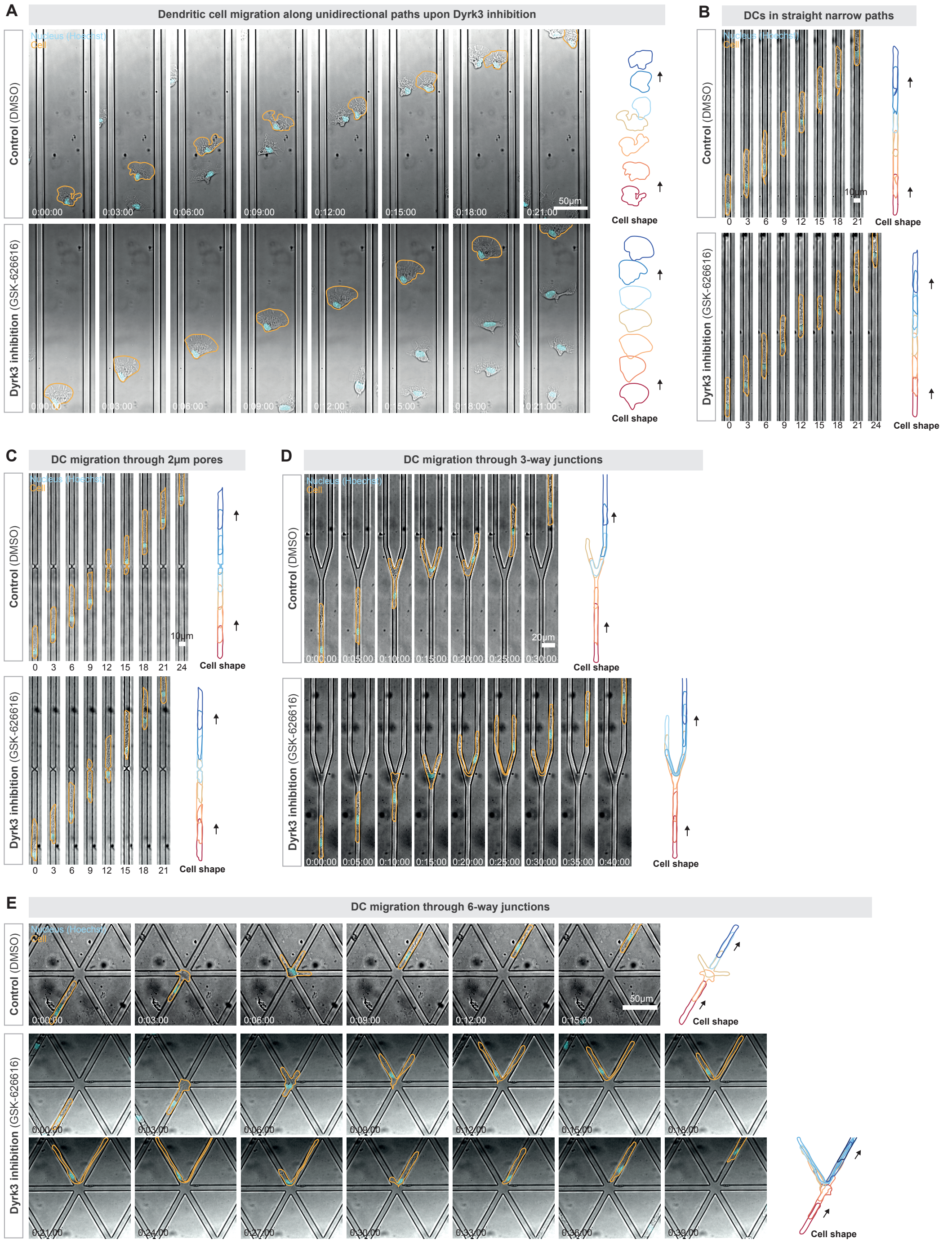


**Fig. S7.**

Dendritic cell migration phenotypes in the presence of mechanically unstable centrosomes (non-functional Dyrk3). **(A)** Representative dendritic cells (DCs) migrating along a unidirectional straight path (wide linear microchannel) in the presence of 5μM GSK-626616 or DMSO (control). See Figure 4E for quantification. **(B)** Representative DCs migrating along a unidirectional straight path (narrow linear microchannel) in the presence of 5μM GSK-626616 or DMSO (control). See Figure 4E for quantification. **(C)** Representative DCs migrating through a 2μm pore in the presence of 5μM GSK-626616 or DMSO (control). See Figure 4F for quantification. **(D)** Representative DCs migrating along a 3-way path junction in the presence of 5μM GSK-626616 or DMSO (control). See Figure 4G for quantification. **(E)** Representative DCs migrating along a 6-way path junction in the presence of 5μM GSK-626616 or DMSO (control). See Figure 4G for quantification. All data show representative cells from at least three independent biological replicates. Time is indicated as h:min:s.

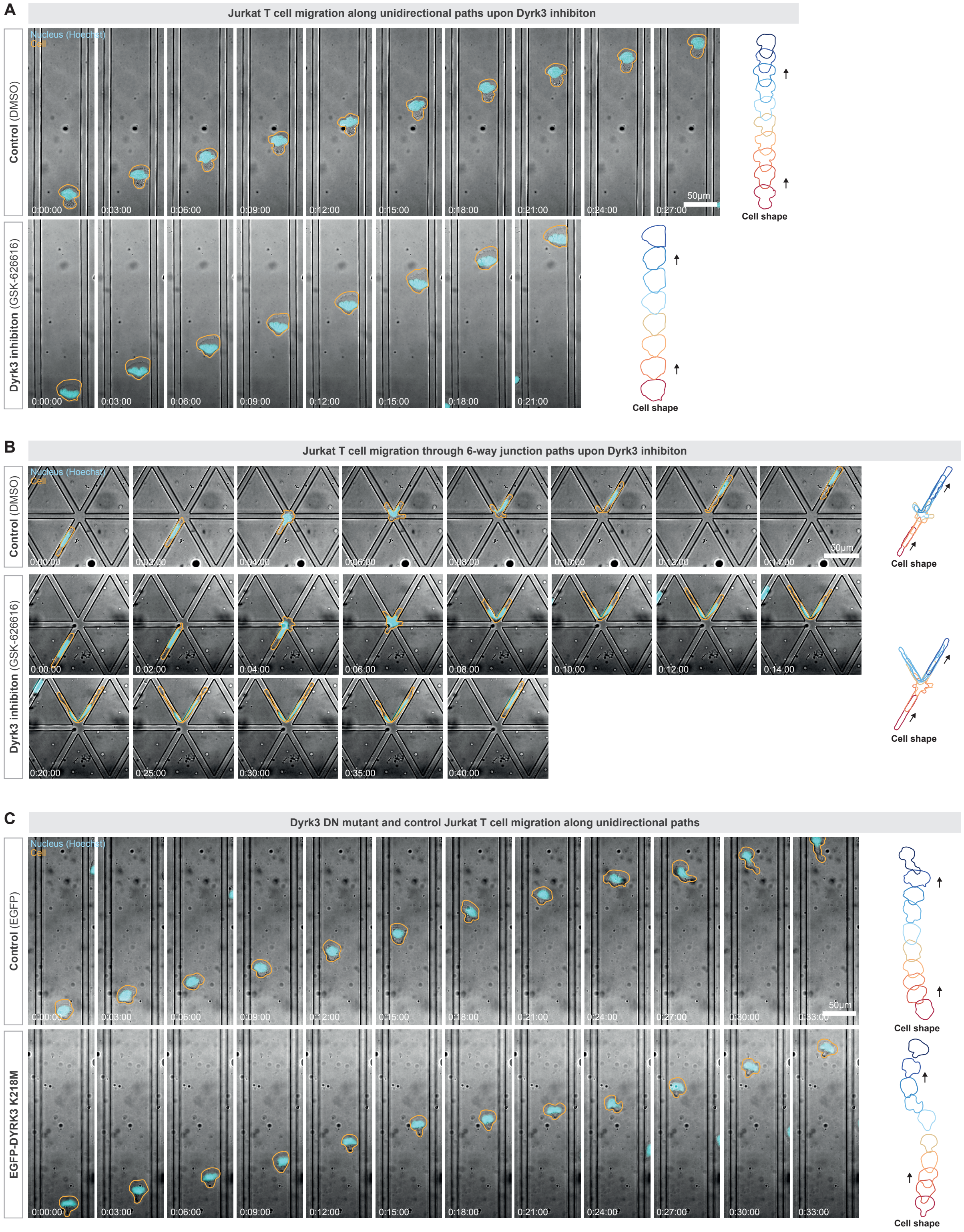


**Fig. S8.**

Jurkat T cell migration phenotypes in the presence of mechanically unstable centrosomes (non-functional Dyrk3). **(A)** Representative Jurkat T cell migrating along a unidirectional straight path (wide linear microchannel) in the presence of 5μM GSK-626616 or DMSO (control). See Figure 4H for quantification. **(B)** Representative Jurkat T cell migrating along a 6-way path junction in the presence of 5μM GSK-626616 or DMSO (control). See Figure 4H for quantification. **(C)** Representative Jurkat T cell expressing EGFP-Dyrk3 K218M or only EGFP while migrating along a unidirectional straight path (wide linear microchannel). See Figure 4J for quantification, and Figure 4I for comparison to migration through path junctions. All data show representative cells from at least three independent biological replicates. Time is indicated as h:min:s.

**Movie S1.**

Mechanical centrosome deformations during cellular navigation. Representative CETN2-GFP expressing dendritic cells migrating along path junctions (1st movie part) and unidirectional paths (2nd movie part) in chemokine gradient (CCL19). The pair of centrioles (CETN2-GFP) is shown in black and the nucleus (Hoechst) is shown in blue. Note the short-range transient separation of the centriole pair (red arrow) in particular during cellular path decisions at path junctions. The movie shows representative cells from at least three independent biological replicates. Time is indicated as h:min:s.

**Movie S2.**

Rendering Dyrk3 non-functional impairs cell migration in three-dimensional collagen matrices. Representative dendritic cells (1st movie part) and Jurkat T cells (2nd and 3rd movie part) migrating in three-dimensional collagen matrices in a chemokine gradient (CCL19), either in the presence of the Dyrk3 inhibitor GSK-626616 (5μM) or DMSO control (1st and 2nd movie part) or in the presence of Dyrk3-K218M-GFP (3rd movie part). The movie shows representative cells from at least three independent biological replicates. Time is indicated as h:min:s.

**Movie S3.**

Centrosome fracturing during cellular navigation upon rendering Dyrk3 non-functional. Representative CETN2-GFP dendritic cells migrating along path junctions (1st movie part) and unidirectional paths (2nd movie part) in the presence of the Dyrk3 inhibitor GSK-626616 (5μM) or DMSO control. The pair of centrioles (CETN2-GFP) is shown in black and the nucleus (Hoechst) is shown in blue. Note the long-range separation of the centriole pair (red arrow) during cellular path decisions in the presence of the Dyrk3 inhibitor GSK-626616. The movie shows representative cells from at least three independent biological replicates. Time is indicated as h:min:s.

**Movie S4.**

Centrosome deformations during cellular squeezing. Representative CETN2-GFP dendritic cells migrating through 2 micrometer pores (1st movie part) and 3 micrometer pores (2nd movie part) in the presence of the Dyrk3 inhibitor GSK-626616 (5μM) or DMSO control. The pair of centrioles (CETN2-GFP) is shown in black; the nucleus (Hoechst) is shown in blue. Note the short-range centrosome deformations during cellular squeezing, which already occurs during translocation through 3 micrometer pores in the presence of the Dyrk3 inhibitor GSK-626616 (5μM). The movie shows representative cells from at least three independent biological replicates. Time is indicated as h:min:s.

**Movie S5.**

Microtubule dynamics in the presence of the Dyrk3 inhibitor GSK-626616. Representative EB3-mCherry (microtubule plus-end binding) expressing dendritic cells in a confining “under-agarose” environment in the presence of the Dyrk3 inhibitor GSK-626616 (5μM) or DMSO control. Growing microtubule plus end tips (EB3-mCherry) are shown in black. The movie shows representative cells from at least three independent biological replicates. Time is indicated as h:min:s.

**Movie S6.**

Dendritic cell migration through path junctions and unidirectional paths in the presence of mechanically unstable centrosomes (non-functional Dyrk3). Representative dendritic cells migrating through wide linear paths (1st movie part), narrow linear paths (2nd movie part), 2 micrometer pores (3rd movie part), 3-way junctions (4th movie part), and 6-way junctions (5th movie part) in the presence of the Dyrk3 inhibitor GSK-626616 (5µM) or DMSO control. The nucleus (Hoechst) is shown in cyan. The movie shows representative cells from at least three independent biological replicates. Time is indicated as h:min:s.

**Movie S7.**

Jurkat T cell migration through path junctions and unidirectional paths in the presence of mechanically unstable centrosomes (non-functional Dyrk3). Representative Jurkat T cells migrating through wide linear paths (1st movie part) or 6-way junctions (2nd movie part) in the presence of the Dyrk3 inhibitor GSK-626616 (5µM) or DMSO control, and representative Jurkat T cells expressing either EGFP (control) or a Dyrk3 mutant (EGFP Dyrk3 K218M) that are migrating through wide linear paths (3rd movie part) or 6-way junctions (4th movie part). The nucleus (Hoechst) is shown in cyan. The movie shows representative cells from at least three independent biological replicates. Time is indicated as h:min:s.
